# A comparison of automated lesion segmentation approaches for chronic stroke T1-weighted MRI data

**DOI:** 10.1101/441451

**Authors:** Kaori L. Ito, Hosung Kim, Sook-Lei Liew

## Abstract

Accurate stroke lesion segmentation is a critical step in the neuroimaging processing pipeline to assess the relationship between post-stroke brain structure, function, and behavior. While many multimodal segmentation algorithms have been developed for acute stroke neuroimaging, few are effective with only a single T1-weighted (T1w) anatomical MRI. This is a critical gap because most stroke rehabilitation research relies on a single T1w MRI for defining the lesion. Although several attempts to automate the segmentation of chronic lesions on single-channel T1w MRI have been made, these approaches have not been systematically evaluated on a large dataset. Here, we performed an exhaustive review of the literature and identified one semi- and three fully automated approaches for segmentation of chronic stroke lesions using T1w MRI within the last ten years: Clusterize, Automated Lesion Identification, Gaussian naïve Bayes lesion detection, and LINDA. We evaluated each method on a large T1w stroke dataset (N=181) using visual and quantitative methods. LINDA was the most computationally expensive approach, but performed best across the three main evaluation metrics (median values: Dice Coefficient=0.50, Hausdorff’s Distance=36.34 mm, and Average Symmetric Surface Distance = 4.97 mm), whereas the Gaussian Bayes method had the highest recall/least false negatives (median=0.80). Segmentation accuracy in all automated methods were influenced by size (small: worst) and stroke territory (brainstem, cerebellum: worst) of the lesion. To facilitate reproducible science, we have made our analysis files publicly available online at https://github.com/npnl/elsa. We hope these findings are informative to future development of T1w lesion segmentation algorithms.

## 1. INTRODUCTION

Despite intensive research and rehabilitation efforts, stroke remains a leading cause of long-term disability world-wide (Mozaffarian et al., 2016). Stroke rehabilitation research aims to understand the relationship between brain, behavior, and recovery following a stroke and to use brain changes after a stroke to predict functional outcomes. Neuroimaging, and in particular, high-resolution T1-weighted (T1w) anatomical MRIs, have been used to examine structural brain changes after stroke. Careful investigation of post-stroke brain anatomy, using techniques such as voxel-lesion symptom mapping (VLSM) or calculation of the overlap percentage between the lesion and critical brain structures, have been useful for relating brain changes to behavioral outcomes (e.g., corticospinal tract lesion load; Bates, Wilson, & Saygin, 2003; Lindenberg et al., 2010; Riley et al., 2011; Zhu, Lindenberg, Alexander, & Schlaug, 2010). However, accurate and precise lesion annotation is necessary to conduct and draw valid clinical inferences from these analyses.

To date, manual lesion tracing by an individual with expertise in neuroanatomy remains the gold standard for lesion segmentation. This procedure is a time and labor-intensive process that requires domain expertise. Consequently, this is not feasible for studies with larger sample sizes, and becomes a limiting factor in large-scale stroke rehabilitation neuroimaging analyses. This is especially problematic for stroke rehabilitation research, as compared to acute stroke research, because there are few stroke segmentation algorithms that can be effectively used with only a T1w MRI for the lesion segmentation. In contrast, there are many more algorithms that have focused on multimodal lesion segmentation.

To clarify the distinction between acute stroke neuroimaging and stroke rehabilitation research, it is useful to note that acute stroke neuroimaging is typically acquired within the first few days after a stroke and used to make clinical decisions about acute stroke treatment. Multimodal MRI sequences are commonly acquired to support this decision-making process, including diffusion weighted imaging, perfusion weighted imaging, T2-FLAIR imaging, and others. A wealth of research attention has focused on developing optimal algorithms for quick lesion segmentation and prediction of gross clinical outcomes using these multimodal sequences.

In contrast, stroke rehabilitation research typically focuses on understanding more fine-grained analyses of functional outcomes following stroke, such as upper or lower limb impairments, aphasia, and more. These studies often acquire research data in stroke participants at a less acute stage, anywhere from several weeks to many years after stroke. Typically, a single high-resolution T1w MRI is acquired to define the lesion, rather than the multimodal clinical imaging acquired at the acute phase. This is typically due to time and financial constraints, as well as the fact that at later chronic stages, swelling and inflammation are no longer expected to be visible. Instead, T1w MRIs are more sensitive to showing cortical necrosis beyond two weeks of stroke onset, and are therefore more suitable for detecting chronic stroke lesions (Allen, Hasso, Handwerker, & Farid, 2012). For rehabilitation research studies, the goal is often to acquire more detailed functional brain scans, such as resting state or task-based fMRI, or detailed anatomical scans such as for diffusion tractography, to understand specific functional and structural patterns associated with post-stroke functional recovery. The downside, however, is that there are fewer lesion segmentation algorithms focused on only T1w MRIs.

In recent years, a handful of automated and semi-automated lesion segmentation approaches have been developed in response to this problem (see literature search results in Supplementary Table 1). Automated segmentation approaches can be divided into two major categories: (1) supervised image classification techniques which use machine learning to train classifiers based on “ground truth” lesion examples (i.e., manually traced lesions), and (2) unsupervised approaches, which use mathematical modeling to first distinguish the lesional tissue characteristics from other tissue types without labeled responses and then separately cluster the voxels belonging to each tissue type. These automated approaches are promising, yet few comparisons between existing T1w lesion segmentation methods have been made, due to (a) the lack of large-sized, publicly available stroke T1w MRI datasets and (b) the intensive labor necessary to manually segment lesions as the benchmark for comparison.

Systematic evaluations of existing algorithms can be useful to identify current best solutions, as well as areas where all algorithms might improve. An excellent example of a comparative evaluation of lesion segmentation algorithms comes from the multimodal, acute neuroimaging world in the form of the annual ischemic stroke lesion segmentation challenge (ISLES challenge; Maier et al., 2017; http://www.isles-challenge.org). ISLES organizers provide a training and testing dataset of multimodal MRI scans acquired from patients with acute stroke. Teams compete to develop algorithms that accurately segment the lesions and upload their algorithms to the ISLES website, after which the ISLES organizers evaluate the lesion segmentations and rank participating research teams based on their performance. The focus of the ISLES challenge is to encourage the development of lesion segmentation methods for acute stroke characterized using multimodal MRI including perfusion MRI, diffusion-weighted imaging (DWI), and fluid-attenuated inversion recovery imaging (FLAIR). Multivariate modeling on these multimodal MRI data increases the sensitivity to early detectable changes after stroke (Allen, Hasso, Handwerker, & Farid, 2012; Baird & Warach, 1998; Yuh et al., 1991).

However, a fair and systematic evaluation of the accuracy of computerized stroke lesion segmentation approaches performed on a large, publicly open dataset of chronic T1w MRIs is needed to facilitate the improvement of their clinical utility in the chronic stroke population. Here, we strived to provide an accurate reflection of the current state of approaches for unimodal T1w chronic stroke lesion segmentation by quantitatively evaluating the performance of existing segmentation algorithms for T1w MRI. We make our analysis files publicly available on our github repository (https://github.com/npnl/elsa) to facilitate reproducibility.

## 2. METHODS

We first performed a review of the literature to identify existing T1w MRI stroke lesion segmentation approaches. We then implemented the identified lesion segmentation approaches, and compared their performance to a ground-truth expert segmentation using various image metrics. Finally, we statistically evaluated how each automated segmentation approach performed against one another.

### 2.1 Literature Search

A computerized search covering the period from April 2007 to April 2017 was conducted on the PubMED online database using the following terms: ((“lesion identification” OR “lesion detection” OR “lesion classification”) OR “lesion” AND (“stroke”[MeSH Terms] OR “brain”[MeSH Terms])) AND “automated”. Studies were limited to those in the English language.

The initial search yielded 189 results. We then excluded articles that were targeted at lesions not caused by stroke (e.g., multiple sclerosis; n=144). Any articles that were not specifically on the topic of lesion segmentation methods were also excluded (n=31). This resulted in 14 remaining results (Supplementary Table 1). Finally, to provide a fair evaluation for algorithms for chronic, T1w stroke MRI, we identified only articles on lesion segmentation approaches that were performed on chronic stroke lesions and have shown to support segmentation on a single T1w (n=6, Table 1). Of these six different lesion segmentation approaches, two were excluded as the software was either no longer available or not supported (S. Shen, personal communication; Shen, Szameitat, & Sterr, 2010; Guo et al., 2015).

**Table 1.**
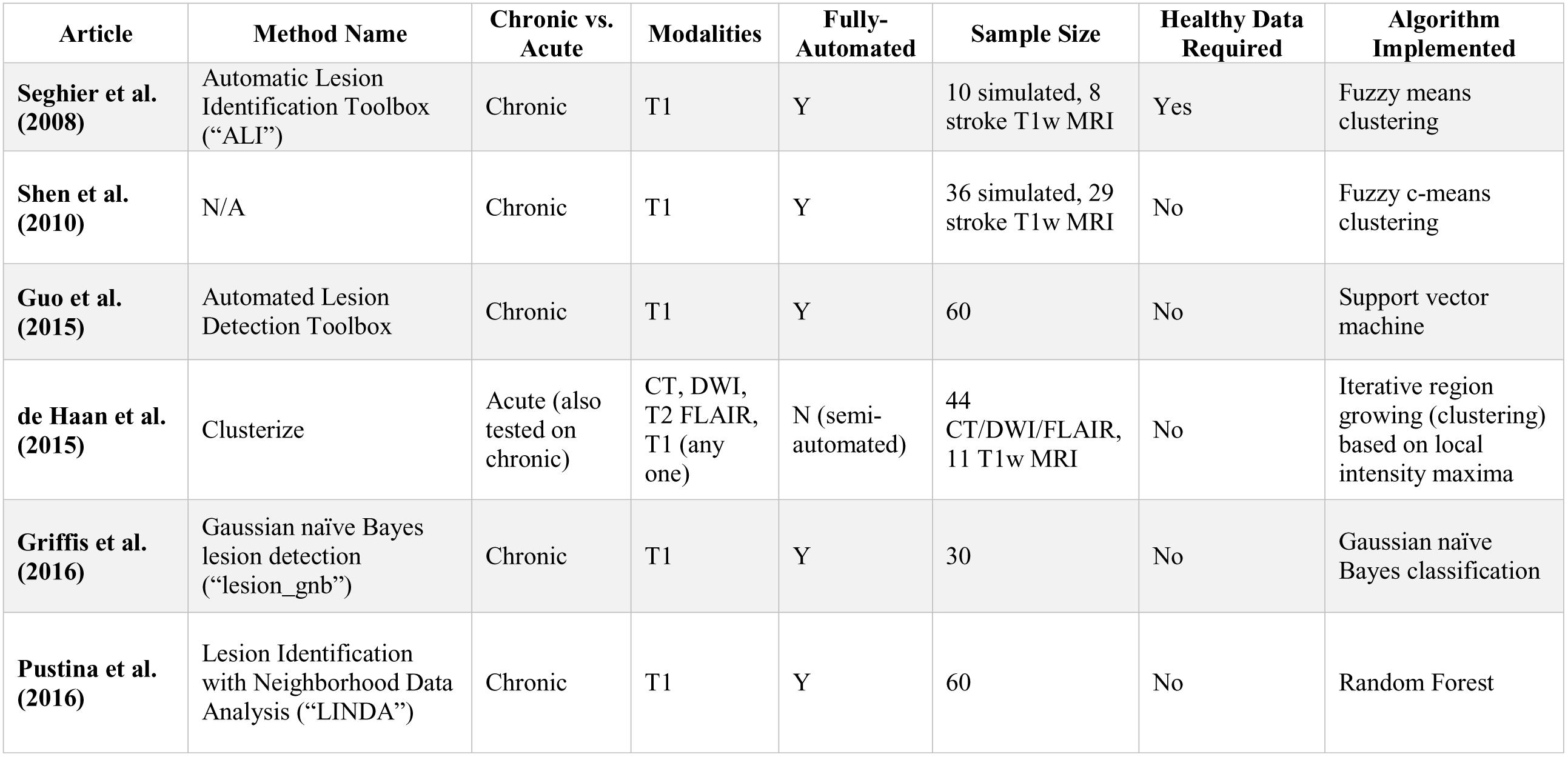
Literature meeting selection criteria. Articles on lesion segmentation approaches for chronic stroke lesions that support a single T1-weighted image segmentation. Two approaches (Shen et al., Guo et al.) were currently unsupported or unavailable and thus excluded from the current evaluations.

| Article | Method Name | Chronic vs. Acute | Modalities | Fully-Automated | Sample Size | Healthy Data Required | Algorithm Implemented |
| --- | --- | --- | --- | --- | --- | --- | --- |
| **Seghier et al. (2008)** | Automatic Lesion Identification Toolbox (“ALI”) | Chronic | T1 | Y | 10 simulated, 8 stroke T1w MRI | Yes | Fuzzy means clustering |
| **Shen et al. (2010)** | N/A | Chronic | T1 | Y | 36 simulated, 29 stroke T1w MRI | No | Fuzzy c-means clustering |
| **Guo et al. (2015)** | Automated Lesion Detection Toolbox | Chronic | T1 | Y | 60 | No | Support vector machine |
| **de Haan et al. (2015)** | Clusterize | Acute (also tested on chronic) | CT, DWI, T2 FLAIR, T1 (any one) | N (semi-automated) | 44 CT/DWI/FLAIR, 11 T1w MRI | No | Iterative region growing (clustering) based on local intensity maxima |
| **Griffis et al. (2016)** | Gaussian naïve Bayes lesion detection (“lesion_gnb”) | Chronic | T1 | Y | 30 | No | Gaussian naïve Bayes classification |
| **Pustina et al. (2016)** | Lesion Identification with Neighborhood Data Analysis (“LINDA”) | Chronic | T1 | Y | 60 | No | Random Forest |

### 2.2 Software Overview

Based on our literature search, we tested both semi- and fully automated approaches to lesion segmentation. In the following sections, a brief overview of each approach is provided.

#### 2.2.1 Semi-Automated Software

We examined one semi-automated approach, Clusterize (Philipp, Groeschel, & Wilke, 2012; de Haan, Clas, Juenger, Wilke, & Karnath, 2015), as it was the only approach that met our literature search criteria. However, we acknowledge that other tools that may be considered ‘semi-automated’, such as MRIcron (http://people.cas.sc.edu/rorden/mricron/index.html) and ITKSnap (Yushkevich et al., 2006), are also publicly available and may be used in lesion segmentation, although they were not developed specifically for stroke lesion segmentation. Both MRICron and ITKSnap provide 3D-growth algorithms to fill an initial area, and require a relatively large amount of manual input to guide the automated segmentation mask, thus making them more comparable to manual lesion segmentation methods. For these reasons, and because they did not meet the literature search criteria, we did not include them in our evaluation.

##### Clusterize Toolbox

The Clusterize approach is a semi-automated approach originally developed to identify demyelination load in metachromatic leukodystrophy using T2-weighted MRIs (Philipp et al., 2012). However, the Clusterize algorithm has been shown to perform comparably on both acute and chronic stroke T1w MRI datasets (de Haan, Clas, Juenger, Wilke, & Karnath, 2015).

The Clusterize approach has an automated preprocessing step followed by a manual cluster selection step. The automated preprocessing step involves identification of the local intensity maxima on each image slice and assignment of each voxel to a single cluster core based on its intensity. This is followed by a manual cluster selection step and an optional freehand correction step to optimize the accuracy of the lesion mask.

#### 2.2.2 Fully Automated Software

Three fully automated approaches resulted from our literature search and were currently available: the automated lesion identification toolbox (ALI), a Gaussian naïve Bayes lesion detection method (lesion_gnb), and lesion identification with neighborhood data analysis (LINDA; Seghier, Ramlackhansingh, Crinion, Leff, & Price, 2008; Griffis, Allendorfer, & Szaflarski, 2016; Pustina et al., 2016).

##### ALI Toolbox

The automated lesion identification (ALI) approach is an *unsupervised* method that performs outlier detection to segment lesions using a fuzzy c-means algorithm (Seghier, Ramlackhansingh, Crinion, Leff, & Price, 2008). The outlier detection procedure includes a voxel-wise comparison between healthy and non-healthy tissue, using a healthy dataset to define the healthy tissue.

##### lesion_gnb

The lesion_gnb approach is a *supervised* method that performs Gaussian naïve Bayes (GNB) classification for the automated delineation of chronic stroke lesions (Griffis et al., 2016). The lesion_gnb approach used 30 training cases to create feature maps encoding information about missing and abnormal tissue, obtained from gray matter (GM), white matter (WM), and cerebrospinal fluid (CSF) prior probability maps (PPM) and tissue probabilistic maps (TPM). The GNB classifier was trained on ground-truth manually delineated lesions as well as these feature maps using a leave-one-out cross-validation approach. The trained GNB classifier is provided by the developers of the lesion_gnb toolbox.

##### LINDA

The LINDA approach is a *supervised* method that relies on feature detection and uses a random forest (RF) algorithm to train and classify lesioned voxels (Pustina et al., 2016). In the LINDA method, features capturing aspects of geometry, subject specific anomalies, and deviation from controls for sixty stroke subjects were fed into a single matrix containing information about a single voxel and its neighboring voxels. The matrix was then used to train the RF algorithm using manually delineated lesions as the ground truth. RF training was repeated two more times with successively hierarchical image resolution. The trained RF classifier is provided by the developers of LINDA.

### 2.3 Data and Implementation of Algorithms

#### 2.3.1 Computational Platform and Software Installation

All computations were performed on a Mac OSX Yosemite operating system with a 3.2 GHz Intel Core i5 processor and 8 GB RAM. To run the ALI, lesion_gnb, and Clusterize toolboxes, we used MATLAB version R2016b and SPM12. For the LINDA toolkit, we used R version 3.3.3, ANTsR version 0.3.1, ANTsRCore version 0.3.7.4, and ITKR version 0.4.12. See Table 2 for more information.

**Table 2.** Processing features of fully automated toolboxes.

|  | Automated Lesion Identification (ALI) | Voxel-based Gaussian Naïve Bayes Classification (lesion_gnb) | Lesion Identification with Neighborhood Data Analysis (LINDA) |
| --- | --- | --- | --- |
| Compatible Operating Systems | Windows, Linux, Mac | Windows, Linux, Mac | Windows 10+, Linux, Mac |
| Platform Dependencies | MATLAB, SPM5+ | MATLAB 2014b+ (Requires Statistics and Machine Learning Toolbox), SPM12+ | R v.3.0+, ANTsR package |
| Year Developed | 2007 | 2015 | 2016 |
| Open Source | No | Yes | Yes |
| Learning Type | Unsupervised | Supervised | Supervised |
| Training Dataset | Requires user to provide segmented healthy training dataset | Provided (trained on 30 left hemisphere stroke subjects) | Provided (trained on 60 left hemisphere stroke subjects) |
| Amenable to left or right hemisphere lesions | Yes | Yes, provided that the user indicates which hemisphere first | No, right hemisphere lesions must be flipped |
| Batch Mode | Yes | Yes (script provided) | Yes (user can write a wrapper script) |
| Template Brain Space | ICBM152 | ICBM152 | Colin 27 template |
| User Defined Parameters | sensitivity (tuning factor), fuzziness index in fuzzy means clustering algorithm, threshold probability and size for the extra class prior | optional smoothing, smoothing kernel, minimum cluster size, implicit masking while smoothing | None |
| Optional Post-Processing Steps | None | Re-segmentation with a tissue prior | None |

#### 2.3.2 Stroke Data

We obtained our stroke dataset from the Anatomical Tracings of Lesions After Stroke (ATLAS) database; (Liew et al., 2018). ATLAS is a public database consisting of 304 T1w anatomical MRIs of individuals with chronic stroke collected from research groups worldwide from the ENIGMA Stroke Recovery Working Group consortium. We excluded any MRIs with non-isometric voxels, as well as any MRIs that highly deviated from the normal range of the standard orientation. We also only included one MRI per individual (no inclusion of longitudinal data). One hundred and eighty-one T1w anatomical MRIs (100 left hemisphere stroke (LHS), 81 right hemisphere stroke (RHS)) from a total of eight different scanners were included in the current analyses. Further information on image acquisition for stroke data can be found in Liew et al., 2018.

#### 2.3.3 Lesion Segmentation

##### Expert Segmentation

The ATLAS database included manually segmented lesion masks created by a team of trained individuals. For the purposes of this evaluation, we included only lesions that were designated as the primary stroke. Each lesion mask was carefully quality-controlled. Briefly, each stroke lesion was segmented using either the coronal or axial view in MRIcron (http://people.cas.sc.edu/rorden/mricron/index.html) with either a mouse, track pad, or a tablet by one of eleven trained individuals, all of whom received detailed training and were compared for inter- and intra-rater reliability on five stroke lesions (inter-rater DC: 0.75±0.18; intra-rater DC: 0.83±0.13; Liew et al., 2018). After lesions were checked for accuracy by a separate tracer, lesion masks were smoothed using a 2mm FWHM kernel in order to smooth jagged edges between slices. For further information on the labeling protocol, see Liew et al., 2018.

##### Semi-automated Segmentation

###### Clusterize

We followed the standard procedure (previously described in 2.2.1) and manually selected clusters as our lesion mask. We did not perform additional manual correction, as this time-consuming process would have made the process analogous to a manual labeling procedure.

##### Automated Segmentations

###### ALI

The ALI method required a dataset of healthy controls to perform outlier detection. As the developers of the algorithm did not provide a healthy control dataset, we fed the algorithm with images of healthy subjects sampled from the Functional Connectome Project (http://fcon_1000.projects.nitrc.org, n=100). The developers of ALI did not recommend an optimal number of healthy controls, but specified that a larger set of healthy controls would better estimate the normal variability in brain structure (M. Seghier; personal communication).

The following automated steps were implemented in the use of the ALI toolbox. All adjustable parameters were kept at their default values. First, segmentation and normalization of both healthy and stroke T1w MRI images were performed in SPM12. For stroke T1w MRIs, the ALI toolbox carried out a modified unified segmentation-normalization algorithm, which included use of an extra lesion tissue class prior (defined as the mean of the standard white matter (WM) and cerebrospinal fluid (CSF) priors) to inform tissue probability maps for the segmentation of GM, WM, and CSF maps. GM and WM segmentations were then smoothed, and outlier detection comparing both GM and WM segmentations between patients and healthy controls was performed using fuzzy means clustering. Finally, the identified GM and WM outliers were combined into a final lesion mask.

###### lesion_gnb

To carry out the lesion_gnb approach, the following steps were implemented: we first specified whether the stroke was on the left or right hemisphere, as the program does not automatically detect the stroke hemisphere. Then, probabilistic tissue segmentation on the stroke T1w MRI was carried out using default parameters in the New Segment tool in SPM12. Tissue segmentations were smoothed with an 8 mm FWHM kernel, and feature maps containing information about missing and abnormal tissue were derived from GM/WM/CSF TPMs. The trained and cross-validated GNB classifier provided by the developers was then used to predict lesion class labels. A minimum cluster size of 100 voxels was specified as the threshold for retention of clusters in the final mask, and the final mask was smoothed using an 8 mm FWHM kernel. The choice of 8 mm FWHM kernel and the cluster size of 100 voxels were based on the default parameters set in the software. Per personal communication, an additional re-segmentation step using the final lesion has been recommended for improving normalization performance and outline precision (J. Griffis, personal communication; Sanjuán et al., 2013). We do not use the additional re-segmentation step for this analysis.

###### LINDA

LINDA requires all strokes to be presented on the left hemisphere. Therefore, as a first step, we mirrored the T1w MRIs for subjects with right hemisphere stroke so that the stroke appeared on the left hemisphere (n=81). The data were preprocessed with two iterations of bias correction and brain extraction, and spatial normalization was performed using Advanced Normalization Tools (ANTs; Avants et al., 2010). Six features (deviation of k-mean segmentation from controls, gradient magnitude, T1 deviation from controls, k-mean segmentation, deviation of T1 asymmetry from controls, and raw T1 volume) were computed from the preprocessed T1w image. These features were fed to the pre-trained RF classifier provided by the developers, and the classifier was then run to detect the lesion at three different image resolutions, where the predicted lesion mask was inversely transformed from the template to the subject’s image space after each iteration to improve prediction accuracy. For right hemisphere lesions, we flipped the images back to the right hemisphere after lesion segmentation.

#### 2.4 Post-Processing of Automated Lesion Masks

As the three fully automated approaches generated lesion masks in their own stereotaxic spaces (i.e., using different templates), all lesion masks were converted back to native space for comparison of automated masks to expert masks. For the ALI and lesion_gnb approaches, lesion masks were inverse transformed using the transformation matrix resulting from SPM. As the LINDA toolbox included a lesion mask output in both native and stereotaxic space, no further processing was necessary for the LINDA approach.

#### 2.5 Segmentation Evaluation

##### 2.5.1 Visual Evaluation

As a first step, we manually inspected the quality of the automated lesion mask outputs. To do so, we used an open-source package (Pipeline for the Analysis of Lesions after Stroke; PALS; http://github.com/NPNL/PALS) to perform a visual evaluation of the automated outputs (Fig. 1; Ito, Kumar, Zavaliangos-Petropulu, Cramer, & Liew, 2018). The Visual QC module in PALS facilitates visual inspection of lesion segmentations by creating HTML pages with screenshots of lesion segmentations overlaid on each subject’s T1w image, and allows for easy flagging of lesion masks that do not pass inspection.

**Figure 1.**
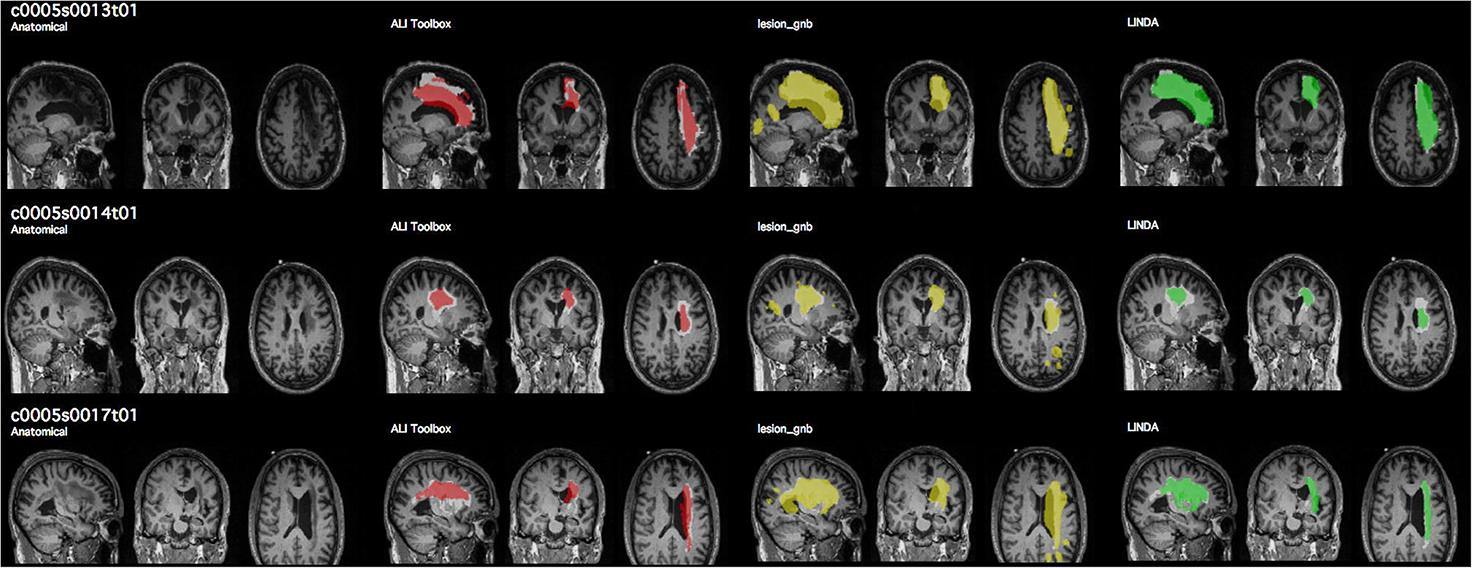
Example of quality control page. Prior to quantitatively evaluating each lesion segmentation performance, we visually assessed the lesion mask for each case. We created a script that automatically output a quality control page (https://github.com/npnl/PALS; Ito et al., 2018) with each automated lesion mask overlaid (red, yellow, green) on the expert segmentation (white). Subject IDs shown in this figure are kept in the same convention as in the ATLAS database.

Given the nature of our multi-site, secondary testing data, we anticipated that there might be cases in which the lesion segmentation algorithms would either 1) produce a lesion mask that has no overlap with the expert segmentation, i.e., misclassify the lesion, or 2) identify no lesioned voxels, creating an empty mask file. For evaluating the performance across approaches, we decided to eliminate a case if any automated approach yielded an empty mask file, as this would not yield any comparable distance metric. Additionally, to provide a fair comparison of approaches, we also decided to eliminate cases in which *all three* automated algorithms produced lesion masks that had no overlap with the expert segmentation, considering them as poor test cases to avoid counting these cases against the segmentation algorithms (Table 3).

**Table 3.**
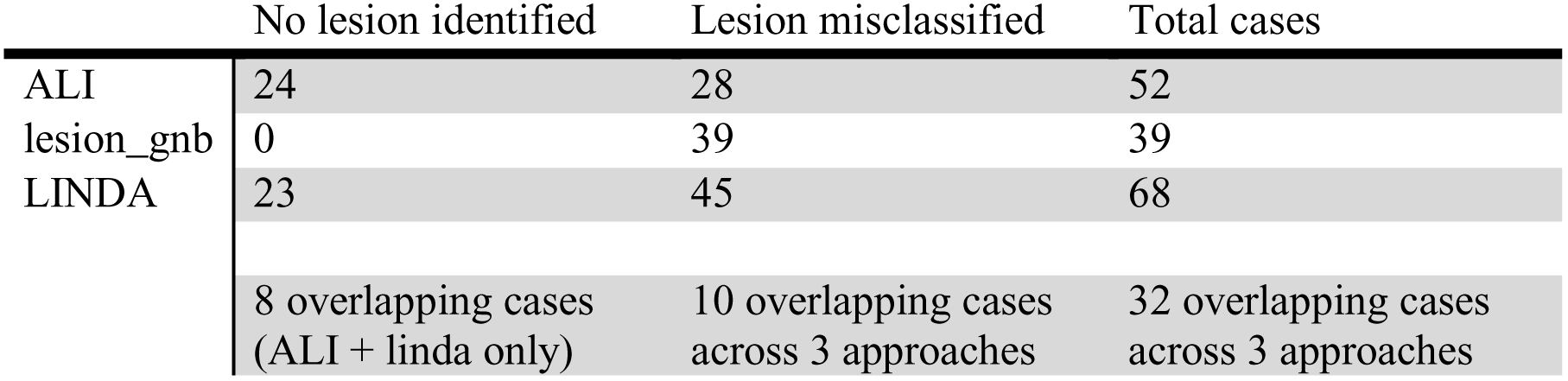
Cases with no lesion mask identified, or lesions misclassified. Cases with no lesion mask identified yielded an empty file (containing only 0 values), and cases in which the lesion was misclassified contained lesioned voxels, but had no voxels overlapping with the expert segmentation. For comparisons between algorithms, we removed cases with no lesions identified (24 ALI + 23 LINDA – 8 overlapping = 39), and removed the 10 cases in which all three algorithms misclassified the lesion.

|  | No lesion identified | Lesion misclassified | Total cases |
| --- | --- | --- | --- |
| ALI | 24 | 28 | 52 |
| lesion_gnb | 0 | 39 | 39 |
| LINDA | 23 | 45 | 68 |
|  | 8 overlapping cases<br>(ALI + linda only) | 10 overlapping cases<br>across 3 approaches | 32 overlapping cases<br>across 3 approaches |

##### 2.5.2 Quantitative Evaluation

###### Evaluation Metrics

To evaluate the performance of each automated lesion segmentation approach compared to the expert segmentation, we implemented the following evaluation metrics: dice similarity coefficient (DC), Hausdorff’s Distance (HD) and Average Symmetric Surface Distance (ASSD). To assess over- and under-segmentation, we also obtained values on precision (also known as positive predictive value) and recall (sensitivity). We additionally calculated the lesion volume to assess whether the automated lesion segmentation approaches detected lesions of similar size to the expert segmentation. Finally, algorithmic efficiency was evaluated by obtaining the computational time for each segmentation approach. Evaluation metrics are described in detail below.

###### Dice Similarity Coefficient

The dice similarity coefficient (DC) is a measure of segmentation accuracy. DC is calculated with the following equation:

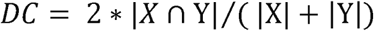

DC ranges from 0 (no overlap) to 1 (complete overlap), and *X* and *Y* represent the voxels in the expert segmentation, and those in the automated segmentation respectively.

###### Hausdorff’s Distance

HD is a measure of the maximum distance between all surface points of two image volumes. It is defined as:

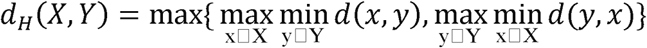

where x and y are points of lesion segmentations X and Y respectively, and d(x,y) is a 3D-matrix consisting of all Euclidean distances between these points. HD is measured in millimeters and a smaller value indicates higher accuracy.

###### Average Symmetric Surface Distance

Average symmetric surface distance (ASSD) is a measure of the average of all Euclidean distances between two image volumes. Given the average surface distance (ASD),

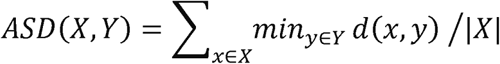

where d(x,y) is a 3-D matrix consisting of the Euclidean distances between the two image volumes X and Y, ASSD is given as:

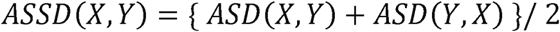

Similar to HD, the ASSD is measured in millimeters, and a smaller value indicates higher accuracy.

###### Precision and Recall

Precision, also called positive predictive value, is the fraction of true positives (that is, overlapping points between the two images) within the automated segmentation. It is defined as:

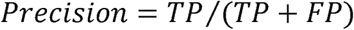

where precision ranges from 0 to 1 (1 indicating optimal precision), and TP are the true positives and FP denotes false positives in the automated segmentation.

Recall, also called sensitivity, is the fraction of true positives (overlapping points between the two images) within the expert segmentation. It is calculated with the equation:

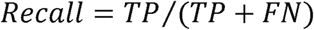

where recall ranges from 0 to 1 (1 indicating optimal recall), and TP denotes true positives and FN (false negative) denotes points that the automated segmentation failed to identify.

#### 2.5.3 Statistical Analyses

All statistical analyses were carried out in R version 3.3.3. To prevent any undue influence of extremely easy or extremely difficult cases, we performed nonparametric analyses to use the ranks of each automated approach to determine whether one approach outperformed another. For fair comparison, our statistical analyses were performed only on the fully automated approaches and did not include the lesion masks created in the Clusterize toolbox, which were driven by a degree of manual input. We report segmentation evaluation metrics for Clusterize in Table 4.

**Table 4.** Descriptive statistics for each approach. Performance rates for each approach; mean ± standard deviation: Dice Coefficient (Dice), Hausdorff’s distance (HD), average symmetric surface distance (ASSD), precision, and recall.

| **Image Metrics**<br>mean ± SD | Clusterize | ALI | lesion_gnb | LINDA |
| --- | --- | --- | --- | --- |
|  | N=152 | N=132 | N=132 | N=132 |
| Dice | 0.23 ± 0.19 | 0.36 ± 0.25 | 0.39 ± 0.23 | 0.45 ± 0.31 |
| HD (*mm*) | 75.00 ± 22.88 | 61.55 ± 28.84 | 58.00 ± 19.73 | 42.07 ± 24.38 |
| ASSD (*mm*) | 13.59 ± 5.85 | 14.38 ± 13.53 | 10.49 ± 6.25 | 12.68 ± 16.49 |
| Precision | 0.16 ± 0.15 | 0.31 ± 0.25 | 0.30 ± 0.20 | 0.50 ± 0.34 |
| Recall | 0.79 ± 0.23 | 0.55 ± 0.31 | 0.69 ± 0.29 | 0.52 ± 0.34 |
| Average Processing Time | 106.43 seconds for automated clustering + 251.75 seconds for manual identification | 396.99 + 247.83 seconds per healthy brain | 246.12 seconds | 3843.66 seconds |

A Friedman test, the nonparametric equivalent of a one-way repeated measures ANOVA, was carried out to examine whether there was a significant difference in the performance among the fully automated segmentation approaches for each evaluation metric (DC, ASSD, HD, precision, and recall). Post hoc analyses with Wilcoxon signed-rank tests were carried out using a Bonferroni correction for multiple comparisons. All Type I error rates were set at α<0.05.

To further evaluate the utility of the automated segmentation approaches on lesion volume, we calculated a Pearson product-moment correlation coefficient for each automated approach to determine the relationship between the lesion volume of the expert segmentation and the lesion volume of the automated segmentation.

#### 2.5.4 Analyses of lesion characteristics in relation to segmentation accuracy

We assessed whether performance of any of the automated lesion segmentation approaches was associated with any particular lesion characteristics, such as stroke territory (cortical, subcortical, brainstem, cerebellar) and lesion size.

Segmentation accuracy was treated as a categorical variable with best and worst performing cases (e.g., based on DC), where DC values were ranked from lowest to highest, and cases below the 25^th^ percentile (Q1) were treated as the worst performing cases, and cases above the 75^th^ percentile (Q3) were treated as best performing cases.

To assess for differences in accuracy in relation to stroke territory *within each approach*, we stratified segmentation accuracy by each automated approach. We then performed a Fisher’s exact test (with a 2 × 4 contingency table including segmentation accuracy and stroke territory variables) on each stratum.

To assess for differences in accuracy *between approaches* within each stroke territory, we stratified segmentation accuracy by each stroke territory. We then performed a Fisher’s exact test (with a 2 × 3 contingency table including segmentation accuracy and segmentation approach variables) on each stratum.

We also created a lesion size variable by transforming lesion volume based on expert segmentations into three categories using the 33^rd^ and 67^th^ percentiles in the dataset of all lesion volumes as cut-off ranges for small, medium, and large lesions. We performed the same Fisher’s exact tests on lesion size as described above.

All statistical tests were adjusted for multiple comparisons using a Bonferroni correction.

## 3. RESULTS

### 3.1 Exploratory Data Analysis

#### 3.1.1 Computational Time

We first examined how long it took for each algorithm to run. For this evaluation, we used a subset of n=100 left hemisphere MRIs (from our total dataset of n=181). This was because additional steps were required for processing right hemisphere lesions in the LINDA toolbox, which would have made it difficult to compare total time across toolboxes. The average times to preprocess an image and detect a lesion for 100 left hemisphere stroke MRIs, in order from fastest to slowest, were as follows: Clusterize (106.43 seconds, but with an additional 251.75 seconds to manually identify each cluster), lesion_gnb (246.12 seconds), ALI (396.99 seconds, but with an additional 247.83 seconds per each healthy brain), and LINDA (3843.66 seconds). Notably, for Clusterize, the manual identification time will vary by user and by lesion. For ALI, which requires healthy brains for comparison, we used 100 healthy brains to match the number of stroke MRIs (see section 2.3.3).

#### 3.1.2 Visual Evaluation

We performed a visual evaluation to assess the quality of the automated lesion masks and ascertain that the lesion masks were correctly transformed back to native space.

For the Clusterize toolbox, there were 152 cases which resulted in a lesion mask, and 29 cases (16.02%) in which no cluster was detected as the lesion mask during manual identification.

All fully automated approaches ran completely and produced a lesion mask file for each case. However, we identified a number of cases where either there were no lesioned voxels that resulted from the automated segmentation, creating an empty file (which we refer to as an empty mask; Table 3), or there was a complete mismatch between the automated and expert segmentation (i.e., all voxels in the automated mask were misclassified as the lesion, which we refer to as a misclassification). This had been anticipated as the algorithms were based on supervised learning with a limited number of their own training data for which lesions in some brain regions (e.g., cerebellum) were not included.

Of the 181 cases, ALI successfully generated 129 lesion masks (71%) with at least a single voxel overlapping with the manual label. Lesion_gnb also detected 142 cases (78%) with at least a single voxel overlap with the manual label, and LINDA detected 113 cases (62%).

There were 24 cases in which ALI produced an empty mask; 23 in which LINDA produced an empty mask (eight of these were the same cases as ALI); and zero in lesion_gnb (in other words, lesion_gnb always created a mask in which it identified what it considered to be lesioned voxels). We subsequently excluded these cases (n=23+24-8=39) from the analysis of the evaluation metrics, as they would not yield any measurable metrics.

In addition, ALI had 28 cases in which the automated segmentation was misclassified (e.g., no overlap between the automated lesion mask and the actual lesion identified by manual segmentation), lesion_gnb had 39 misclassified cases, and LINDA had 45 cases (Table 3). Ten of these were the same cases across all three approaches, and we removed these ten misclassified cases from evaluation, treating them as poor test cases. Hence, after exclusion of 39 cases that had empty masks and 10 misclassified cases, 132 total cases remained in our quantitative evaluation.

For a discussion on the implications and possible reasons for misclassification and failed lesion detection, see section 4.4).

### 3.2 Quantitative Evaluation

The performance of each fully automated toolbox was evaluated across the following metrics: Dice Similarity Coefficient, Hausdorff’s Distance, Average Symmetric Surface Distance, Precision, and Recall (Fig. 2, 3). A summary of the findings, in which we ranked the toolboxes based on their relative performance to one another, can be found in Table 5.

**Figure 2.**
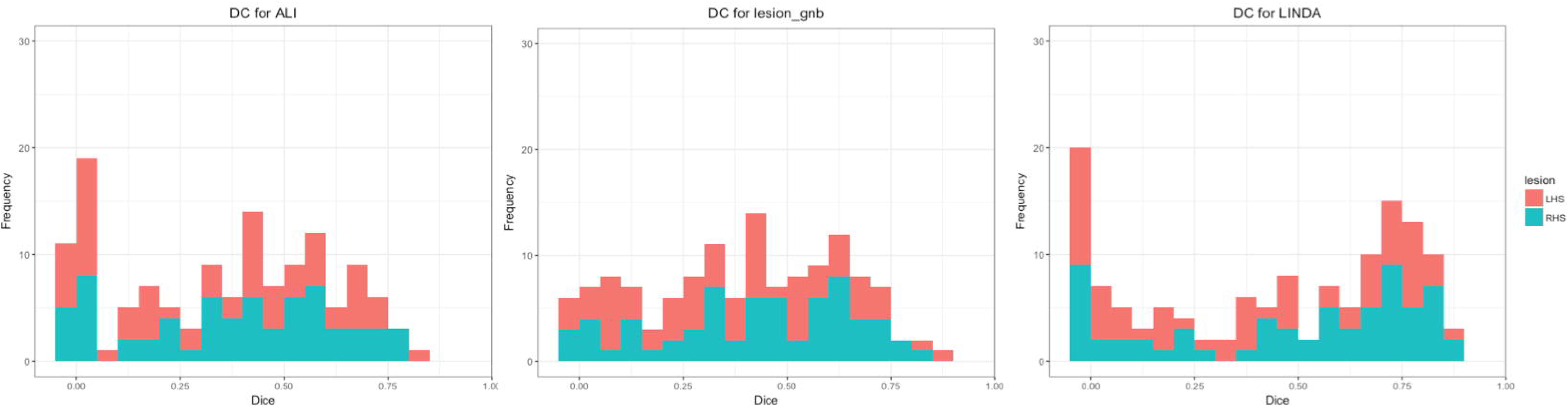
Distribution of Dice Similarity Coefficient values for automated approaches. Histograms of all dice similarity coefficient values (N=132) for each automated lesion detection approach; left hemisphere stroke (LHS) in orange; right hemisphere stroke (RHS) in turquoise.

**Figure 3.**
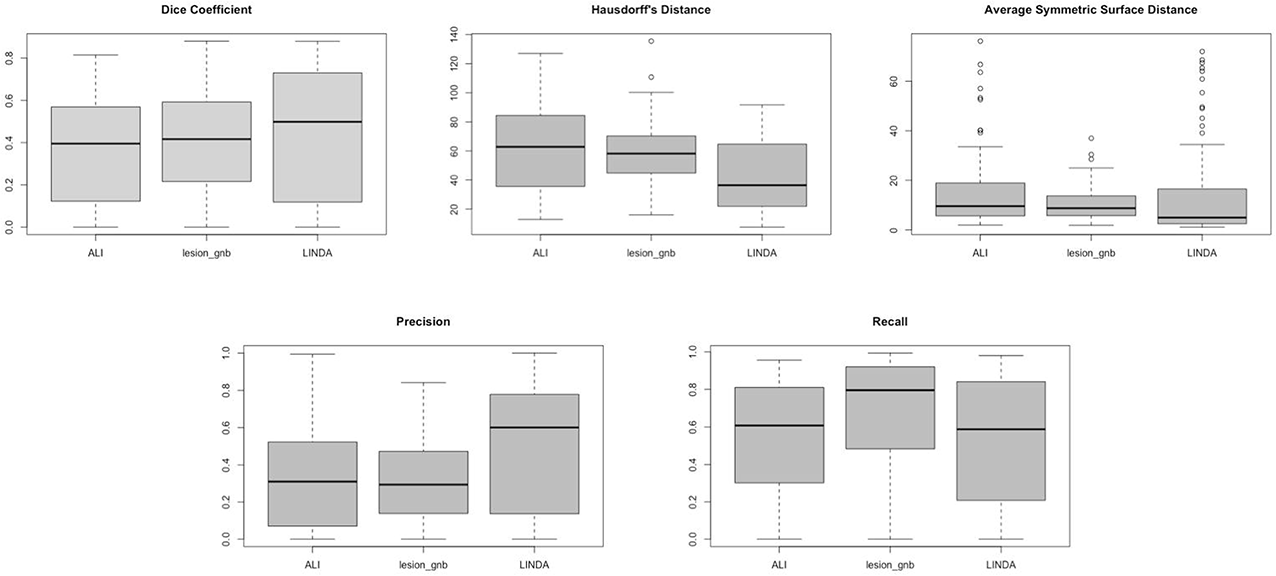
Boxplots of evaluation metrics for automated approaches. For Dice Coefficient, Precision, and Recall, range is from 0-1, where 0=worst and 1=best; Hausdorff’s Distance and Average Symmetric Surface Distance are measured in millimeters, and smaller values indicate better performance.

**Table 5.**
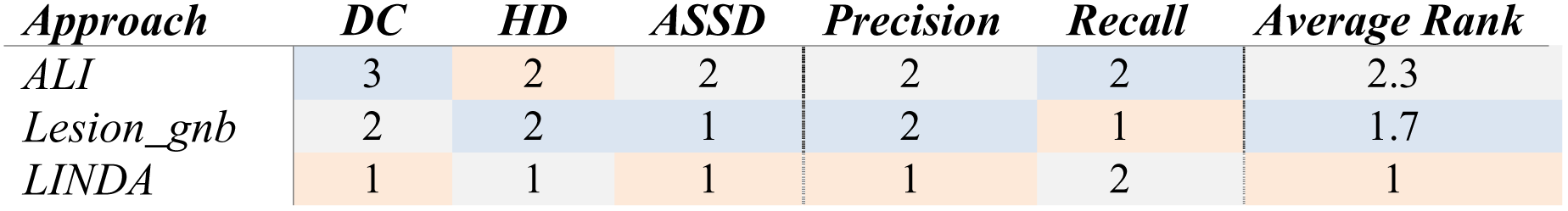
Ranks of segmentation performance on each evaluation metric. Performance ranked by median values for each approach (if the approach significant performed better than the others); dice coefficient (DC), Hausdorff’s distance (HD), average symmetric surface distance (ASSD). Average rank only takes the mean of DC, HD, and ASSD as precision and recall are reflected in the dice coefficient.

#### 3.2.1 Dice Similarity Coefficient

Using a Friedman test, we found a statistically significant difference in DC among the three fully automated lesion segmentation approaches, χ^2^(2)= 27.10, p<0.0001; corrected using the Bonferroni adjustment which was applied to all the following tests. Median (IQR) DC values for ALI, lesion_gnb, and LINDA approaches were 0.40 (0.00 to 0.81), 0.42 (0.00 to 0.88), and 0.50 (0.00 to 0.88), respectively. Post hoc analyses using Wilcoxon signed-rank tests on DC showed that LINDA outperformed lesion_gnb and ALI (sum of positive ranks, lesion_gnb: V=5359, p=0.01; ALI: V=1944, p<0.0001), and lesion_gnb outperformed ALI (V=2601, p<0.0001).

#### 3.2.2 Hausdorff’s Distance

A statistically significant difference in ranks for HD was found among the three fully automated segmentation approaches, χ^2^(2)=43.09, p<0.0001. Median (IQR) HD values for ALI, lesion_gnb, and LINDA are as follows: 62.79 mm (12.81 to 127.10), 58.19 mm (16.06 to 135.50), and 36.34 mm (7.55 to 91.75), where smaller values indicate better performance. Wilcoxon signed-rank tests showed that LINDA performed better than ALI and lesion_gnb (ALI: V=7156, p<0.0001; lesion_gnb: V=1915, p<0.0001). There were no significant differences between ALI and lesion_gnb (V=5085, p=0.34).

#### 3.2.3 Average Symmetric Surface Distance

We also found a statistically significant difference in ASSD among the three fully automated segmentation approaches, χ^2^(2)=42.97, p<0.0001. Median (IQR) ASSD values for ALI, lesion_gnb, and LINDA approaches were 9.58 mm (1.94 to 76.11), 8.75 mm (1.88 to 36.94), and 4.97 (1.11 to 71.95), respectively. Again, smaller values indicate better performance. Pairwise comparisons showed that LINDA and lesion_gnb both performed better than ALI (LINDA: V=6832, p<0.0001; lesion_gnb: V=5990, p=0.0008), but there were no significance differences between LINDA and lesion_gnb (V=3766, p=0.47).

#### 3.2.4 Precision and Recall

We found a statistically significant difference in median precision among the three fully automated approaches, χ^2^(2)=41.59, p<0.0001. Median (IQR) precision values for ALI, lesion_gnb, and LINDA approaches were 0.31 (0.00 to 0.99), 0.29 (0.00 to 0.84), and 0.60 (0.00 to 1.00), respectively. Here, higher values indicate better performance. Wilcoxon signed-rank tests showed that LINDA had higher precision rates than both ALI (V=1423, p<0.0001) and lesion_gnb (V=6961, p<0.0001), and there were no significant differences between ALI and lesion_gnb (V=3998, p=1.00).

We also found a statistically significant difference in recall among the three fully automated lesion segmentation approaches, χ^2^(2)= 97.86, p<0.0001. Median (IQR) recall values for ALI, lesion_gnb, and LINDA approaches were 0.61 (0.00 to 0.96), 0.80 (0.00 to 0.99), and 0.59 (0.00 to 0.98), respectively, again with higher values indicating better performance. We found that lesion_gnb performed better than both LINDA (V=1084, p<0.0001), and ALI (V=1171, p<0.0001), and there were no significant differences between LINDA and ALI (V=4479, p=0.55).

### 3.3 Volume Correlation

For each of the fully automated segmentation approaches, we also examined the relationship between the lesion volumes of the automated segmentations and the lesion volumes of the expert segmentations across the 132 cases. To account for outliers, we removed observations for which the Cook’s distance was greater than 1 after fitting a simple linear regression. Two observations were removed for LINDA, and one was removed for ALI and lesion_gnb each. After removing outliers, we calculated the Pearson correlation coefficient, and found a statistically significant positive correlation between the lesion volumes of the expert segmentations and the lesion volumes of each of the automated segmentations: ALI (r=0.75, p<0.0001), lesion_gnb (r=0.90, p<0.0001), and LINDA (r=0.84, p<0.0001). This suggests that the lesions that were automatically detected were of similar volume to those of expert segmentations.

### 3.4 Lesion Characteristics of Highest and Lowest Performing Cases

We then examined whether there were specific characteristics of the lesion that led to good or bad algorithm performance. Lesion characteristics (hemisphere of the lesion, stroke territory, and lesion size) for both best and worst performing cases based on DC values (top and bottom 25%) are provided in Table 6. For these comparisons, we included all 181 cases, treating cases which did not identify any lesioned voxels as having DC=0, as we wanted to examine what may have contributed to the cases in which lesion masks were misclassified or unidentified. Overall, across all lesion toolboxes, we found the worst algorithm performance for lesions brainstem or cerebellar regions, and for lesions that were small. Unsurprisingly, we found the best algorithm performance in cortical regions and for large lesions.

**Table 6.**
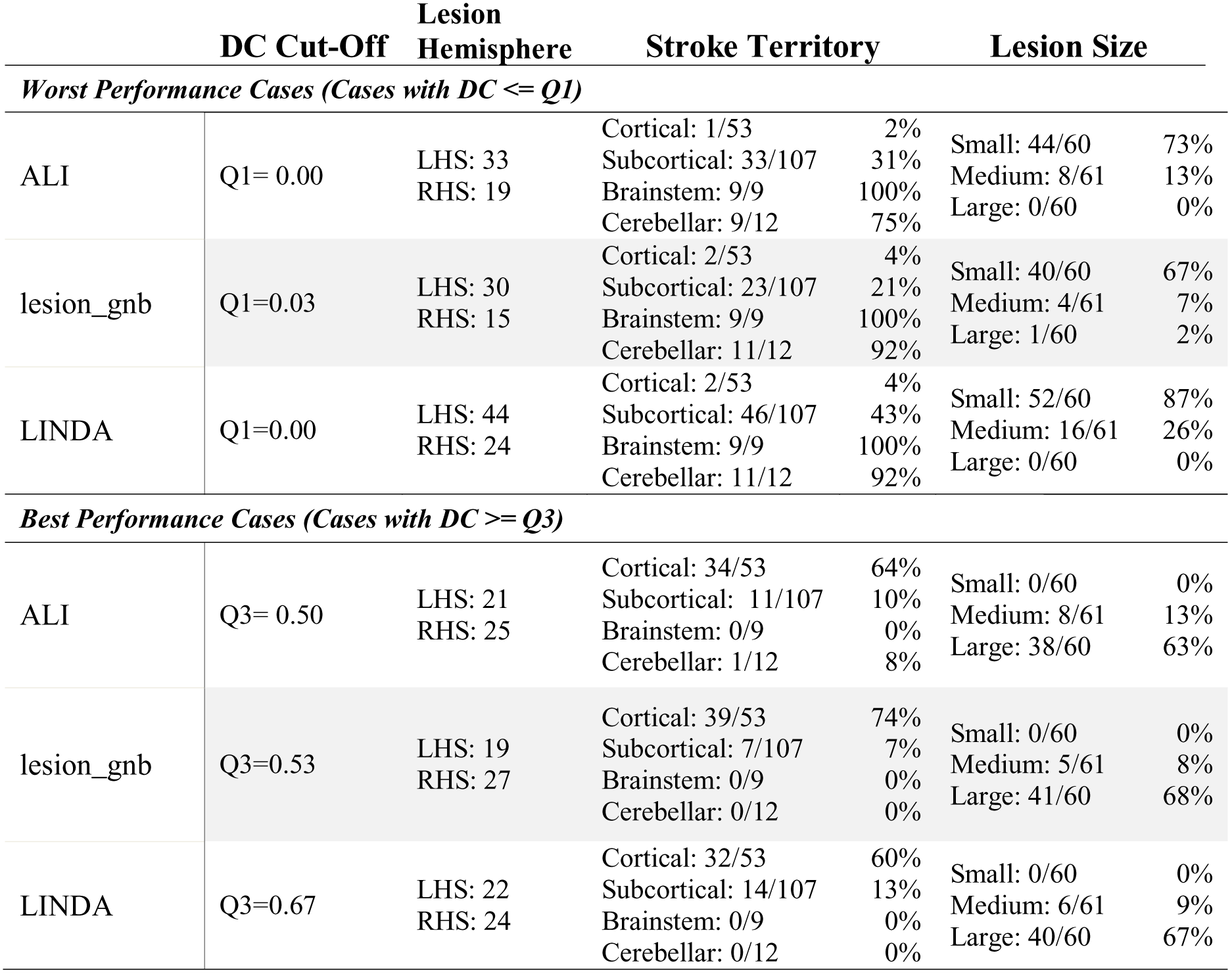
Lesion characteristics based on Dice Coefficient value. Lesion characteristics of best and worst performing cases based on DC values (top and bottom 25%). All 181 cases are included.

#### 3.4.1 Analyses of Cases by Stroke Territory

For ALI, fisher’s exact test showed a significant difference in segmentation accuracy (low performing (DC < Q1) vs. high performing (DC > Q3)) among the stroke territories (cortical, subcortical, brainstem, cerebellar; p<0.0001). Pairwise comparisons showed significant differences between frequency of cortical lesions and all other types of lesions (subcortical, brainstem, and cerebellar; p<0.0001), where cortical lesions performed better than the rest. For lesion_gnb, fisher’s exact test with Bonferroni correction showed a significant difference in segmentation accuracy (low performing (DC < Q1) vs. high performing (DC > Q3)) among the stroke territories (cortical, subcortical, brainstem, cerebellar) (p<0.0001). Again, pairwise comparisons further showed significant differences between frequency of cortical lesions and all other types of lesions (p<0.0001), with cortical lesions performing better than the rest. For LINDA, we also found a significant difference in segmentation accuracy among the stroke territories (p<0.0001). Again, pairwise comparisons showed that significant differences in frequency of cortical lesions to other types of lesions, with cortical lesions performing best (p<0.0001).

Between the automated approaches, we did not find significant differences in frequency of high versus low performing cases for any category of stroke territory (p>0.3).

#### 3.4.2 Analyses of Cases by Lesion Size

For ALI, fisher’s exact test showed a significant difference in segmentation accuracy among the different lesion sizes (p <0.0001). Between low performing and high performing cases, significant differences were found between frequency of small and mid-sized lesions, small and large lesions, and mid-sized and large lesions (p<0.0002), where overall, low performance cases had more small lesions, and high performing cases had more large lesions.

For lesion_gnb, we also found a significant difference in segmentation accuracy among the lesion sizes (p<0.0001). Again, between low and high performing cases, significant differences were found between frequency of small and mid-sized lesions, small and large lesions, and medium and large lesions (p<0.02), where low performance cases had more small lesions and high performing cases had more large lesions.

Similarly for LINDA, we found a difference in segmentation accuracy among the different lesion sizes (p<0.0001). Again, between low and high performing cases, significant differences were found between frequency of small and mid-sized lesions, small and large lesions, and medium and large lesions (p<0.004), where low performance cases had more small lesions and high performing cases had more large lesions.

Between automated approaches, we did not find any significant differences in frequency of high performing cases for lesion size categories (p>0.7).

### 3.5 Misclassified Cases

Finally, we analyzed the cases in which the automated algorithms detected lesions, but did not correctly identify any voxels overlapping with the expert segmentation—in other words, there was an automated lesion mask was created, but the dice coefficient yielded 0. As noted earlier, this occurred in 28 cases for ALI, 39 cases for lesion_gnb, and 45 cases in LINDA. For each of these cases, we quantified the minimum distance (d_min_) between the edge of the expert segmentation with the edge of the automated segmentation with the following equation:

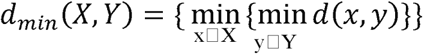

where x and y are points of lesion segmentations X and Y respectively, and d(x,y) is a 3D-matrix consisting of all Euclidean distances between these points. Minimum distance is measured in millimeters and a smaller value indicates higher accuracy. We examined this minimum distance measure to better understand, when lesion masks missed the lesion completely, if they were at least close to the actual lesioned territory, or far off.

Average d_min_ are as follows: ALI: 36.10 ± 21.72 mm (range: 2.24 - 93.25 mm); lesion_gnb: 19.31 ± 13.73 mm (range: 1.41 – 42.91 mm); and LINDA: 29.67 ± 19.82 mm (range: 1.00 – 83.36 mm). A density plot of minimum distances to the manual segmentation is shown in Fig 4. As the plot shows, masks from lesion_gnb were closest to the lesion, followed by ALI and then LINDA. However, due to the low precision of lesion_gnb (i.e., high false positive rate), it is likely that lesion_gnb creates multiple false positive labels, some of which may have been in closer proximity to the true lesion.

**Figure 4.**
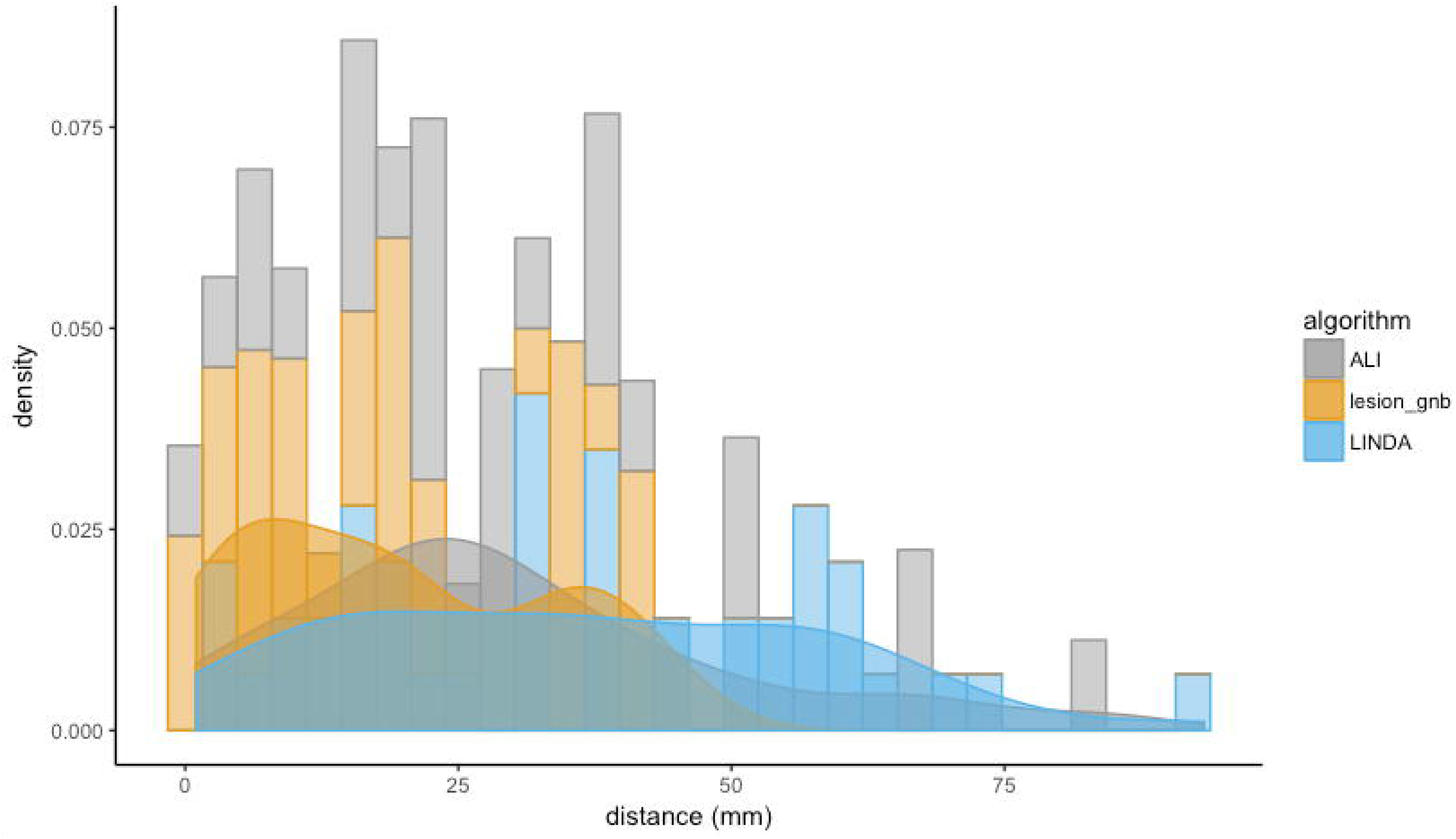
Density plot of minimum distances to manual segmentations for lesions with no overlap. As the plot shows, lesion masks from lesion_gnb were closest to the lesion, followed by ALI, and then LINDA. However, due to the low precision of lesion_gnb (median = 0.29), it is likely that lesion_gnb creates multiple false positive clusters, some of which end up being closer to the manual segmentation.

## 4. DISCUSSION

In the present paper, we systematically evaluated the performance of existing stroke lesion segmentation approaches for chronic T1w MRI on a large common dataset. We tested the accuracy of automated lesion segmentations against a ground-truth expert segmentation by employing a number of image evaluation metrics, including the dice similarity coefficient, Hausdorff’s distance, and average symmetric surface distance. Additionally, we computed precision and recall, measured computational time, calculated the Pearson’s correlation between lesion volumes of expert to automated approaches, and investigated lesion characteristics of highest- and lowest- performing cases. Finally, we analyzed the distance between expert segmentations and automated segmentations that did not yield any voxels that overlapped with the expert segmentation. Overall, we found that LINDA performed the best out of the fully automated lesion segmentation methods. In addition, all methods performed the worst on small lesions, as well as lesions in the brainstem and cerebellum. These finds provide implications for how to improve existing lesion segmentation algorithms for T1-weighted MRIs.

### 4.1 Fully Automated Software

Our findings showed that each of the fully automated approaches resulted in different patterns in the various evaluation metrics used in the current study, indicating that each approach had its own benefits and drawbacks. Specifically, we found that lesion_gnb yielded the least number of cases (in fact, 0) in which no lesion mask was detected, compared to both LINDA and ALI. However, LINDA consistently performed the best out of the evaluation metrics (dice coefficient, Hausdorff’s distance, average symmetric surface distance).

A closer look at precision and recall provides insight to the results obtained from the DC and distance metrics: LINDA resulted in higher precision (positive predictive value) rates than both ALI and lesion_gnb, while recall values were highest in lesion_gnb and similar in ALI and LINDA. Moreover, LINDA had roughly equivalent median precision and recall values (0.60 and 0.59, respectively), whereas both ALI and lesion_gnb had relatively better recall compared to precision (ALI precision: 0.31, recall: 0.61; lesion_gnb precision: 0.29, recall: 0.80). This suggests that both the lesion_gnb and ALI approaches tended to over-segment lesions (high false positives). These findings were confirmed by our visual evaluation.

Our findings suggest that LINDA was the approach that consistently performed best across all metrics—but only when it successfully identified a lesion. However, LINDA was also the most computationally expensive approach: the average time to process a single image on LINDA took roughly sixteen times as long as lesion_gnb, and six times as long as ALI. Additionally, in cases in which automated segmentations were misclassified, the misclassified lesion was in closer proximity to the expert segmentation for lesion_gnb and ALI as compared to LINDA. It is also important to note that both ALI and lesion_gnb have adjustable parameters that may have potentially improved performance when they are adapted to the current test data. As mentioned above, in order to systematically evaluate performance without bias from expert feedback, however, we implemented the approaches with their default settings. Hence, we recommend that the user use his or her discretion to weigh the benefits and drawbacks of each approach as we have presented here, and tailor the settings to his or her specific dataset.

### 4.2 Lesion Detection Algorithms and Methodology

The three fully automated segmentation approaches evaluated here implemented distinct methodologies in their approach to lesion segmentation. ALI used an unsupervised approach with fuzzy means clustering to detect outliers in gray and white matter segmentations, lesion_gnb used a supervised naïve Bayesian classification algorithm to estimate the probability of a lesion class, and LINDA used a supervised random forest approach with a multi-resolution framework to classify voxels and their neighbors as lesional tissue. We expected that the supervised learning algorithms would have higher performance than an unsupervised approach, given that supervised approaches are trained with ground-truth lesions. Indeed, we found that both LINDA and lesion_gnb had higher values than ALI on the dice coefficient. Yet this was not consistently the case for the distance metrics (ASSD and HD). However, these approaches implemented different image processing tools (e.g., ANTs, SPM) that include various preprocessing steps, such as brain extraction, registration and tissue classification. It is likely that performance variability in these preprocessing steps may have had downstream effects on lesion segmentation accuracy. Manual quality control and inspection of preprocessing steps could enhance the lesion segmentation process, and would ideally be a part of any neuroimaging analysis pipeline.

Previously reported results from the developers of each automated algorithm provide a useful tool for comparison and evaluation of the results we obtained from the current study. Our DC values were approximately 0.20 - 0.24 lower than those reported by the developers in their original papers (ALI: original = 0.64, ATLAS = 0.40; lesion_gnb: 0.66, 0.42; LINDA: 0.70, 0.50; Griffis, Allendorfer, & Szaflarski, 2016; Pustina et al., 2016; Seghier et al., 2008). There are several likely explanations for this. First, each of these automated algorithms was originally tested on single site, single scanner-acquired data. This makes these algorithms vulnerable to over-fitting to their own data. In particular, the supervised methods (LINDA, lesion_gnb) were dependent on machine learning classifiers that were pre-trained using data acquired from a single scanner from the original study. Here, we implemented a large data evaluation and tested each pre-trained algorithm on multi-site data. Not surprisingly, we found a significant drop in segmentation accuracy, as variability in machine characteristics was likely not addressed using the initial training-set. Second, to provide an equal comparison across toolboxes, we implemented the fully automated approaches as they were, without modifications to the parameters selected using the original training dataset. We also kept built-in preprocessing steps prior to lesion detection. While performance of the approaches may have been improved by fine-tuning the default parameters to our dataset, the current results obtained without modifications provide valuable baseline information to researchers and clinicians who may be interested in using any of the tested algorithms and wish to bypass the time- and computationally-intensive training procedure.

### 4.3 Small Subcortical, Brainstem, or Cerebellar Lesions Perform Worst

We also assessed whether automated lesion segmentation performance was related to specific lesion characteristics. Overall, we found that the fully automated approaches were less likely to detect small lesions, with half of all total small lesion cases (30/60) failing to be detected by all three fully automated approaches. This is consistent with the literature, which has shown that automated and semi-automated approaches for T1w lesion segmentation to be biased for detection of large lesions (Wilke et al., 2011; Griffis et al., 2016). Users of these algorithms should thus manually inspect lesion segmentation quality, and pay specific attention to small lesions. However, as these are typically the fastest to manually segment, they should also be the fastest to correct. Using an automated segmentation algorithm may therefore still save considerable time, even with manual inspection and corrections for smaller lesions.

Regarding lesion location, fully automated approaches displayed significantly higher segmentation accuracy on cortical lesions than subcortical, brainstem, and cerebellar lesions. As brainstem and cerebellar strokes occur less frequently, brainstem and cerebellar lesions were likely not included in the original training set for the automated algorithms (Chua & Kong, 1996; Datar & Rabinstein, 2014; Kase, Norrving, Levine, & Babikian, 1993; Teasell, Foley, Doherty, & Finestone, 2002). Moreover, features implemented in the algorithms to classify lesions (e.g., hemispheric asymmetry) may not be sensitive to brainstem or cerebellar strokes. Finally, subcortical, brainstem, and cerebellar lesions are often smaller than cortical lesions, suggesting a potential additive effect on accuracy. Users with datasets containing multiple brainstem or cerebellar strokes may need to re-train the algorithm with a training dataset that contains more of these types of lesions to increase algorithm sensitivity.

### 4.4 Dropped Cases

In our evaluation, we found that there was a fairly sizeable number of cases in which lesions were not detected for the ALI and LINDA algorithms. One potential explanation for this is that the thresholds that were used to classify lesioned versus nonlesioned voxels was too high. We had opted to use default values in implementing the algorithms, since we wanted to be systematic in our evaluation, and because ALI had user-adjustable parameters but LINDA did not. However, this meant that the thresholds that were used might not have been optimally tuned for this dataset. Although here the focus was to fairly evaluate the various algorithms using a common set of parameters, an actual user who is trying to generate lesion masks for his or her data could try to better optimize a specific method for a specific dataset.

Additionally, we found that there were a number of cases in which lesions were misclassified. That is, cases where the automated algorithm generated a lesion mask that did not overlap at all with the lesion identified from the manual segmentations. Of misclassified cases, lesion_gnb produced lesion masks that were closest to the true, expert segmentation. However, we note that this may in fact be due to the increased false positive rate of the lesion_gnb algorithm. Notably, across all of the segmentation algorithms, most of these cases were small and brainstem/cerebellar lesions, which may not be reflective of the cases that were used in the training datasets of the algorithms (for a comparison of lesion volumes in our test cases and those used in training datasets, see Supplementary Table 2). Finally, the number of cases with sub-optimal segmentations may also have been inflated in due to our use of secondary, combined multi-site data, which would also increase the variability in lesion characteristics, and cause it to deviate from those used in training datasets.

### 4.5 Semi-Automated Software

We tested one semi-automated software, the Clusterize toolbox, for lesion segmentation. Notably, this toolbox was designed for use with manual input and corrections. We only performed the initial manual step of cluster selection (identifying the lesioned region), but did not perform the subsequent manual correction. The automated preprocessing plus manual cluster selection resulted in a relatively low DC value (M=0.18, IQR: 0.06, 0.37), but a fairly high recall value (sensitivity; M=0.89, IQR: 0.71, 0.96). The high recall was likely driven by the manual selection of clusters: due to expert feedback, a cluster corresponding to the true lesion was accurately selected for most cases. However, Clusterize tended to overestimate the lesioned region, which led to lower precision in the lesion segmentation. In particular, we found that the cluster corresponding to the true lesion often additionally included the ventricle as part of the lesion when the lesion was adjacent to the ventricle. These lower precision values may also have been partly driven by the fact that Clusterize was originally designed for the detection of a different type of the brain lesion (i.e., metachromatic leukodystrophy) on T2-weighted images, suggesting a need for additional feature modeling or parameter optimization. The creation of a mask for the ventricles and exclusion of any voxels within that mask could also enhance this method.

## 5. CONCLUSION

Our systematic evaluation facilitates and informs future use and development of automated approaches. Notably, we found that the supervised algorithms performed best, but there was a high failure rate across all approaches. We also found systematic differences in segmentation accuracy depending on stroke territory and size. Based on these findings, we recommend two primary areas for improvement in the future development of automated lesion detection algorithms: (1) that algorithms be trained on larger and more diverse datasets, allowing for inter-scanner variability from multi-site, multi-scanner data, and (2) that prior knowledge about lesion size and territory be integrated into algorithms to increase segmentation performance. For clinicians and researchers who wish to use currently available lesion detection approaches, we suggest that they carefully select an automated lesion detection approach most suitable for their purposes and perform a thorough visual inspection of the automated segmentations to ensure the accuracy of each mask. We strongly recommend manual quality control following any of these approaches. By facilitating and informing the use and development of automated segmentation approaches, we hope that this systematic review will advance the discovery of clinically meaningful findings about stroke recovery.

## ACKNOWLEDGEMENTS

We would like to thank Allie Schmiesing for her contribution to the literature search and performing lesion segmentations on the Clusterize Toolbox and Jennifer Chan for performing segmentations on the Clusterize Toolbox and her comments on this paper. We would also like to thank all of the ATLAS contributors and contributors to the Functional Connectome Project healthy dataset.

## SOURCES OF FUNDING

This work was supported by the National Institutes of Health NCMRR [grant number 1K01HD091283 to S-LL]. HK was supported by Baxter Foundation Fellowship Award.

## CONFLICTS OF INTEREST

We have no conflicts of interest to declare.

